# Differential momentary reports of stress and affect associated with alcohol consumption in middle-aged versus younger adults

**DOI:** 10.1101/2021.10.25.465757

**Authors:** Hope Peterson, W. Jack Rejeski, Jason Fanning, Stephen W. Porges, Keri J. Heilman, Paul J. Laurienti, Lise Gauvin

**Author notes:** Funding: NIAAA P50 AA026117, NIAAA P01 AA02099, NIAAA T32 AA007565.

## Abstract

**Objective:** Stress is a motivator to consume alcohol, a well-documented relapse risk, and is known to differentially affect biological and psychological processes as people age. Because alcohol consumption is known to decrease stress and increase affect, this study examined differences in ratings of stress and affect across the day in middle-aged versus younger adults who regularly consumed alcohol.

**Methods:** A sample of males and females including younger (n=17) and middle-aged (n=18) drinkers was studied across two experimental periods: a 3-day period of usual drinking and a 3-day period of abstinence from alcohol. We also measured resting levels of respiratory sinus arrhythmia (RSA_rest_), since it is a well-documented biomarker of stress and known to decrease with age. Ecological momentary assessment (EMA) ratings across periods of normal drinking and abstinence were modeled using hierarchical regression to assess differences in stress and affect throughout days of abstinence and normal drinking between the two age groups.

**Results:** As anticipated, middle-aged participants had lower RSA_rest_ than those who were younger. Our analyses showed that middle-aged adults experienced a significant reduction in stress following drinking while no such effect was observed in the younger adults. Although the middle-aged adults showed overall lower stress, generally they also expressed higher affect than younger adults.

**Conclusions:** Despite the mitigating role of alcohol on stress in the middle-aged group and the fact that they had higher affect than the younger adults, their lower levels of RSA_rest_ and higher daily reports of stress could pose a risk for chronic alcohol consumption.

## Introduction

The effects of stress are modulated by age. Although most studies show that younger and older adults typically engage in the same approaches to coping with stress (Aldwin, 1991), stress in early life can affect nervous and immune system responses to stress later in life (Graham et al., 2006; Porges, 1995, 2011a), and older adults more often show immunological impairment in response to stress in comparison to younger adults (Graham, et al., 2006). Additionally, a growing body of evidence documents decreasing mental health and increasing stress among middle-aged adults, compared to those of the same age in previous decades (Almeida et al., 2020; Infurna et al., 2021; Kirsch et al., 2019). Overall well-being shows a U-shaped distribution by age, with middle-aged adults having lower well-being that younger and older adults (Barry & Jenkins, 2007; Lykken & Tellegen, 1996; McEwin & Wingfield, 2010; McKnight-Eily et al., 2021; Miyawaki et al., 2020; Nes et al., 2006). Stress effects also vary by sex, with pronounced and multi-faceted sex differences in stress reactivity in the central nervous system that span neurochemical, behavioral, and cognitive responsiveness (Bowman et al., 2003; Kajantie & Phillips, 2006; Luine, 2009; Merz & Wolf, 2016). Stress is an important factor known to increase alcohol relapse risk (Brown et al., 1995; Noone et al., 1999; Sinha, 2007, 2011, 2012), but stress is also a major public health concern on its own: one third of adults report increasing stress over the past year with an increasing proportion reporting extreme stress (Babor, 2002; Babor & Higgins-Biddle, 2000). This increase in stress is additionally concerning due to the relationship between stress, motivation to drink, and dysfunctional drinking behaviors (Breese et al., 2006; Miller et al., 1974; Seo et al., 2011; Uhart & Wand, 2008).

It is well documented that metrics of heart rate variability such as resting respiratory sinus arrhythmia (RSA_rest_) are inversely related to age, even in the context of healthy aging (Barantke et al., 2008; Dywan et al., 2008; Hrushesky et al., 1984; Parati et al., 1997; Schwartz et al., 1991) and are a well-documented biomarker of stress (Porges, 1995). The amplitude of respiratory sinus arrhythmia (RSA), which is directly proportional to high frequency heart rate variability (HFHRV) (Billman, 2011), falls approximately 10% with each decade of life (Hrushesky, et al., 1984). This age effect contributes approximately 15% to individual differences in RSA_rest_ (Byrne et al., 1996). Although there are also known sex differences in HFHRV, with females generally showing higher HFHRV than males (Beauchaine et al., 2008; Snieder et al., 2012; Vidal-Ribas et al., 2017; Wu et al., 2010), these differences seem most prevalent in younger adults, with decreasing sex difference with increasing age (Beauchaine, et al., 2008; Fagard, 2001; Snieder, et al., 2012). Interestingly, alcohol cravings, chronic consumption, and acute consumption are all associated with decreases in HFHRV (Ganesha et al., 2013; Pietila et al., 2018; Quintana et al., 2013; Quintana, 2013) and excessive alcohol consumption can result in autonomic dysfunction, marked by depressed HFHRV (Julian et al., 2020). In males classified as alcoholics, decreased RSA_rest_ is associated with poor mood and increased stress (Romero-Martinez et al., 2019).

It is well known that middle-aged adults are now increasing their regular alcohol consumption (GBD, 2018), and data from the National Institute of Aging suggest that heavy drinking can make health problems worse as adults grow older (Breuninger et al., 2020). Alcohol consumption is known to decrease stress, and although those with alcohol use disorders (AUD) drink for a variety of reasons, many ingest alcohol to reduce daily stress (Barros Guinle & Sinha, 2020; Park et al., 2004; Peltier et al., 2019; Pohorecky, 1991; Steptoe et al., 1996). Moreover, alcohol consumption is known to acutely increase affect and improve mood particularly among social alcohol consumers (Barros Guinle & Sinha, 2020; Hussong et al., 2001; Steptoe, et al., 1996). With these data in the mind, the current study examined daily stress and affective responses of younger and middle-aged adults who regularly consumed alcohol in response to brief periods of normal drinking and abstinence. We also assessed RSA_rest_ as a biomarker of stress in these participants. We hypothesized that the middle-aged adults would exhibit higher levels of stress across the day and poorer affect than younger adults, effects that would be corroborated by lower levels of RSA_rest_ and mitigated following episodes of drinking.

## Methods

Study participants (n=35; 20 females) included individuals between the ages of 24-60 years, who were right-handed, and consumed alcohol habitually. This study sought to examine moderate to heavy drinkers and as such required participants whose drinking was above NIAAA low risk standards (NIAAA, 2011): ≥7 drinks per week for females and ≥14 drinks per week for males. Additionally, we included only participants that drank on at least 50% of days and had maintained a similar drinking pattern for at least 3 years. Consumption patterns were assessed using the Timeline Followback, a self-report estimate of alcohol consumption over the past 90 days (Vakili et al., 2008). Exclusion criteria included use of other substances (screened with a saliva drug test, with additional limits of 500 mg of caffeine daily, and a nicotine limit of the equivalent of 1.5 packs of cigarettes daily), an alcohol use disorder (AUD) diagnosis or self-reported problems with alcohol, a history of withdrawal symptoms, binge drinking (NIAAA, 2011) more than once monthly on average, regular morning drinking or “ eye openers” (alcohol consumed before noon), diagnosis of neurological disease or psychiatric conditions or history of head injury.

During an initial screening visit, participants completed a battery of self-report questionnaires probing stress (Cohen et al., 1983), anxiety (Julian, 2011), and mindfulness (Walach et al., 2006); as well as the Childhood Trauma Questionnaire (Bernstein et al., 1997; Bernstein et al., 2003; Scher et al., 2001), the Alcohol Craving Experience survey (Statham et al., 2011), the Multidimensional Assessment of Interoceptive Awareness (Mehling et al., 2018), and the Body Perception Questionnaire short form (Cabrera et al., 2018). Heart rate data were also collected in the initial screening visit, using a 3-electrode electrocardiogram (ECG) setup through a Biopac Bionomadix system. Participants were asked to relax and sit comfortably in a chair without speaking or moving for 5 minutes.

Following the screening visit, participants were issued an Apple iPhone device and given one-on-one instruction on a web-based EMA application created for the study. Participants completed surveys across two experimental sessions: a 3-day period of their normal drinking routine, and 3 days of complete alcohol abstinence (occurring in a sex-stratified crossover randomization). Across both sessions, participants were asked to complete EMA surveys upon waking, before beginning a drinking episode, after finishing a drinking episode, before going to sleep, and approximately every two hours throughout the day when prompted between 9 AM and 10 PM. Surveys asked participants to rate their current stress on an unmarked scale ranging from “ not at all” to “ extremely,” and their current affect (how bad or good they feel) on an unmarked scale ranging from “ bad” to “ good”. Scales that participants saw included no numerical values and were designed to slide continuously to allow participants to provide a depiction of their current state. The data were collected using integer scales from 1 (not at all) to 1,000 (extremely) for stress levels and -5 (bad) to 5 (good) for affect.

CardioEdit software (Brain-Body Center, University of Illinois at Chicago, 2007) was used to visually inspect and edit off-line heart rate data. CardioBatch Plus software (Brain-Body Center for Psychophysiology and Bioengineering, University of North Carolina at Chapel Hill, 2016) was used to calculate heart rate and RSA_rest_ from the ECG data, consistent with previously developed procedures (Porges, 1985). CardioBatch Plus quantified the amplitude of RSA_rest_ using age-specific parameters that are sensitive to the maturational shifts in the frequency of spontaneous breathing. This method, when applied to adults, includes: (1) timing sequential R-R intervals to the nearest millisecond; (2) producing time-based data by resampling the sequential R-R intervals into 500 ms intervals; (3) detrending the time-based series with a 21-point cubic moving polynomial stepped through the data to create a smoothed template, then subtracting the template from the original time-based series to generate a detrended residual series; (4) bandpass filtering the detrended time series to extract the variance in the heart period pattern associated with spontaneous breathing in adults (0.12 – 0.40 Hz); and (5) transforming the variance estimates with a natural logarithm to normalize the distribution of RSA estimates (Rinolo & Porges, 1997). RSA was quantified during each sequential 30 sec epoch and the averages within each condition were used in the data analysis.

To examine age and its associated factors, we divided the sample into middle-aged and younger groups based on a median split of age (Johnson, 1967; Lee, 1981), a value that mirrored the established lower bound for middle-aged adults. We then compared groups based on scores of their autonomic function (i.e. RSA_rest_). Hierarchical or nested mixed modeling was then used to assess variance in EMA stress and affect responses prior to and following drinking and as a function of time of day. Survey responses were concatenated and tallied and all responses on normal drinking days were stratified as pre-drinking and post-drinking, using two dummy variables. Responses on abstinence days thus became the reference. On all surveys, time of day was recorded in 24-hour notation, and for analyses, time was centered at 15:00 (3 PM). Responses collected following midnight were assigned times greater than 24:00 to classify them to the day on which they occurred (e.g. 25:00 for 1 AM). When creating the statistical model, the centered time was squared to operationalize quadratic trends to account for potential nonlinear variations in ratings across the day (Mayhugh, Laurienti, et al., 2018; Mayhugh, Rejeski, et al., 2018). This allowed for modeling of linear increases or decreases throughout the day, as well as potential curvilinear patterns of responding. The multivariate modeling determined the influence of within and between subjects variation (Raudenbush, 2001). This statistical model explored within-day variations in ratings and across drinking states (Mayhugh, Laurienti, et al., 2018; Mayhugh, Rejeski, et al., 2018). All EMA analyses were conducted using HLM 8 software (Raudenbush et al., 2019).

The hierarchical mixed-model results presented here were generated assessing untransformed stress and affect ratings. However, because patterns of EMA responding were not normally distributed for stress, several additional analyses were performed to serve as sensitivity analyses. These models were also run controlling for sex, modeling stress responses as a Poisson distribution, calculating z-scores to examine person-centered standardized ratings and running stress and affect the models, and finally by creating a dichotomous variable indicating participants scoring in the top quintile of responses compared to all others and modeling. The multi-level equations generated with the statistical model are included in the supplemental materials.

## Results

A total of 35 participants (20 females) completed the full study. Overall, participants averaged 41.5 years of age (median = 43 years). The full sample of participants had been consuming alcohol for nearly 25 years, consuming an average of 17 drinks per week or 3 drinks per day. No significant differences were found between male and female participants when comparing age, RSA_rest_, or total number of years drinking, but males reported drinking 23 drinks per week or 3.7 drinks per day, while females reported consuming 13 drinks per week or 2.4 drinks per day (*t* = 5.22, *p* < 0.001). All participant characteristics are included in Table 1.

**Table 1.** Participant characteristics for male and female participants and as a function of sex and age-group membership.

|  | **Overall**<br>**N = 35** |  | **Middle-**<br>**aged group**<br>**N = 18** | **Younger**<br>**adults**<br>**N = 17** |  | **Males**<br>**N = 15** | **Females**<br>**N = 20** |
| --- | --- | --- | --- | --- | --- | --- | --- |
| **Age** | 41.54<br>(12.18) |  | 52.28 (5.49) | 30.18 (4.04) |  | 39.5<br>(10.15) | 43.55<br>(13.41) |
| **RSA<sub>rest</sub>** | 5.87 (1.44) |  | 5.13 (1.05) | 6.66 (1.39) |  | 5.82 (1.30) | 5.95 (1.56) |
| **Total years drinking** | 24.7 (11.8) |  | 35.06 (5.83) | 13.88 (4.74) |  | 21.9 (9.2) | 26.9 (13.2) |
| **Average consumption**<br>**per week** | 17 (5) |  | 17.34 (7.24) | 18.88 (8.14) |  | 23 (6) | 13 (5) |
| **Average consumption**<br>**per day** | 2.9 (0.9) |  | 2.48 (1.03) | 2.69 (1.16) |  | 3.7 (1.2) | 2.4 (0.7) |

As designed, the middle-aged group (n=18) had a higher average age [M (SD) = 52.28 (5.49)] than the younger adults [(n=17), 30.18 (4.04)] and, as expected, had levels of lower RSA_rest_ [5.13 (1.05) versus 6.66 (1.39), respectively]. There was no significant difference in the respective proportions of women and men between the groups (*t* = -1.161, *p* = 0.254), with 12 females and 6 males (66% female) in the middle-aged group and 8 females and 9 males in the younger group (47% female). There was no overlap in the age range between the groups (younger: 24-36; middle-aged: 43-59), although there was overlap in values of RSA_rest_ [middle-aged =2.50 (6.89); younger = 4.67(8.17)]. There was no significant difference in the number of drinks consumed on average between the two groups (*t* = 0.597, *p* = -.554), but middle-aged group members had been consuming alcohol for significantly more years than younger group members [35.06 (5.83) versus 13.88 (4.74); t = -11.751, p < 0.001]. This effect was likely driven by the difference in age between groups, as age and total number of years drinking were highly correlated (middle-aged, *r* = 0.96, *p* < 0.001; younger, *r* = 0.92, *p* < 0.001; full sample *r* = 0.98, *p* < 0.001).

Hierarchical mixed modeling of EMA patterns (n = 1598) across the two groups showed significant differences in ratings of stress (Table 2) and affect (Table 3). First, patterns of stress ratings across the day followed a significant curvilinear shape (γ = -0.57, *p* = 0.010), showing stress scores decreased following the centered military time of 15:00. Middle-aged adults showed a significantly greater decrease in afternoon stress ratings than younger adults (γ = -0.74, *p* = 0.016). Only middle-aged adults showed a significant decrease in ratings of stress following alcohol consumption (γ = -88.98, *p* < 0.001). A visual representation of the predicted values is included in Figure 1. Second, patterns of affect ratings across the day showed a significant difference in overall intercept (γ = 1.23, *p* = 0.010) and significant differences between groups in both the linear (increasing before 15:00; γ = -0.04, *p* = 0.002) and curvilinear slopes (decreasing after 15:00; γ = -0.007, *p* = 0.013). Pre-drinking affect scores were not significantly different between the groups, and neither group showed a significant change in affect following drinking.

**Table 2.** Results of multilevel modeling analyses examining the impact of time of day and occurrence of drinking on ecological momentary assessment ratings of stress as a function of group membership. Full model equations can be found in the supplemental materials.

|  |  |  |  |
| --- | --- | --- | --- |
| **Intercept** |  |  |  |
| Overall | $\gamma_{00} = 365.23$ | SE = 37.09 | $p < 0.001$ |
| Middle-aged group | $\gamma_{01} = -65.74$ | SE = 51.71 | $p = 0.213$ |
| **Time of Day – Slope** |  |  |  |
| Overall | $\gamma_{10} = -2.09$ | SE = 1.29 | $p = 0.105$ |
| Middle-aged group | $\gamma_{11} = -0.85$ | SE = 1.78 | $p = 0.635$ |
| **Time of Day –**<br>**Curvilinear** |  |  |  |
| Overall | $\gamma_{20} = -0.57$ | SE = 0.22 | $p = 0.010$ |
| Middle-aged group | $\gamma_{21} = -0.74$ | SE = 0.31 | $p = 0.016$ |
| **Pre-Drinking Slope** |  |  |  |
| Overall | $\gamma_{30} = -25.02$ | SE = 14.02 | $p = 0.075$ |
| Middle-aged group | $\gamma_{31} = -28.20$ | SE = 19.46 | $p = 0.147$ |
| **Post-Drinking Slope** |  |  |  |
| Overall | $\gamma_{40} = -15.18$ | SE = 16.36 | $p = 0.354$ |
| Middle-aged group | $\gamma_{41} = -88.98$ | SE = 22.75 | $p < 0.001$ |

\* Statistically significant findings are highlighted for ease of reading.

**Table 3.**
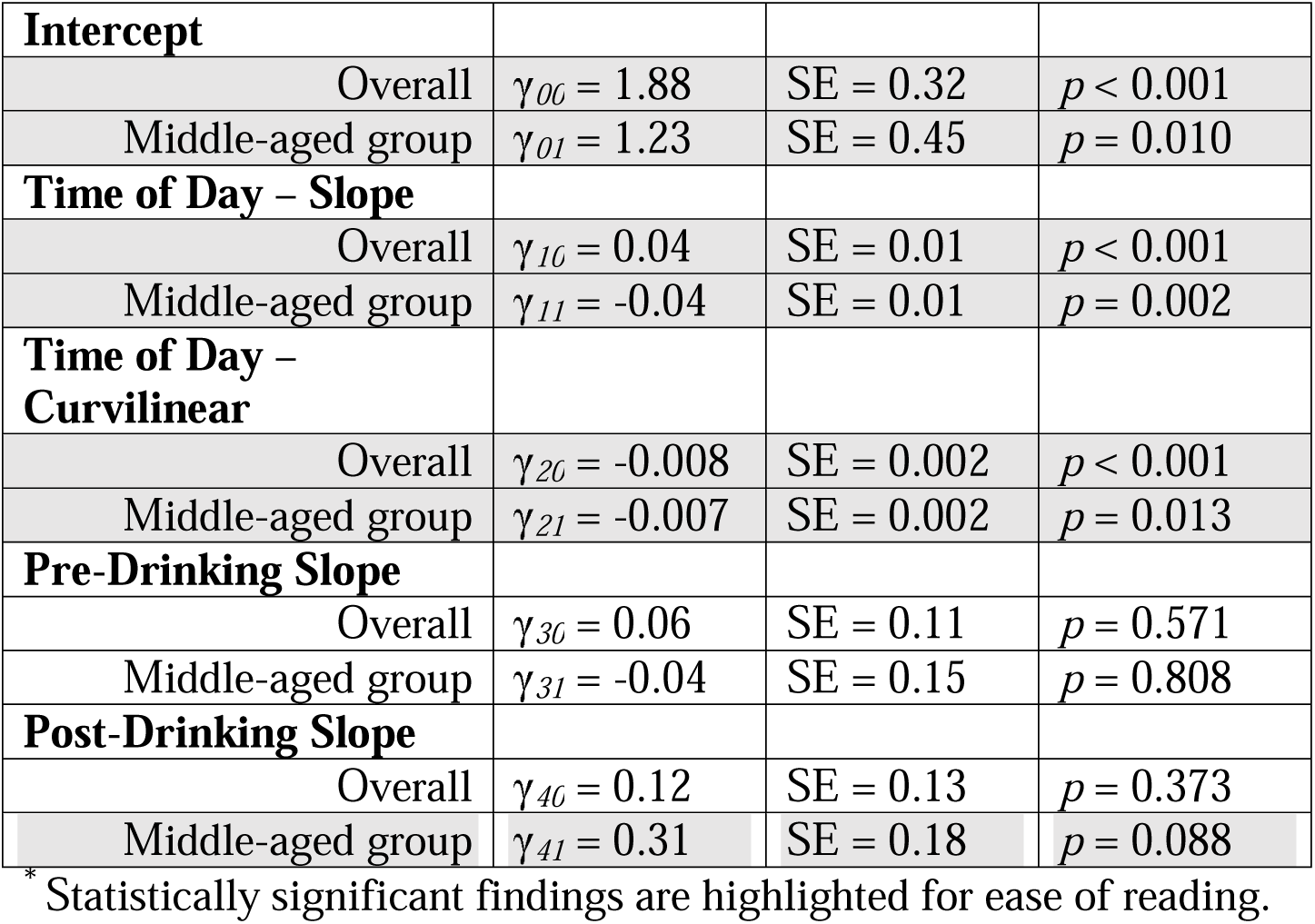
Results of multilevel modeling analyses examining the impact of time of day and occurrence of drinking on ecological momentary assessment ratings of affect as a function of group membership. Full model equations can be found in the supplemental materials.

|  |  |  |  |
| --- | --- | --- | --- |
| **Intercept** |  |  |  |
| Overall | $\gamma_{00} = 1.88$ | SE = 0.32 | $p < 0.001$ |
| Middle-aged group | $\gamma_{01} = 1.23$ | SE = 0.45 | $p = 0.010$ |
| **Time of Day – Slope** |  |  |  |
| Overall | $\gamma_{10} = 0.04$ | SE = 0.01 | $p < 0.001$ |
| Middle-aged group | $\gamma_{11} = -0.04$ | SE = 0.01 | $p = 0.002$ |
| **Time of Day – Curvilinear** |  |  |  |
| Overall | $\gamma_{20} = -0.008$ | SE = 0.002 | $p < 0.001$ |
| Middle-aged group | $\gamma_{21} = -0.007$ | SE = 0.002 | $p = 0.013$ |
| **Pre-Drinking Slope** |  |  |  |
| Overall | $\gamma_{30} = 0.06$ | SE = 0.11 | $p = 0.571$ |
| Middle-aged group | $\gamma_{31} = -0.04$ | SE = 0.15 | $p = 0.808$ |
| **Post-Drinking Slope** |  |  |  |
| Overall | $\gamma_{40} = 0.12$ | SE = 0.13 | $p = 0.373$ |
| Middle-aged group | $\gamma_{41} = 0.31$ | SE = 0.18 | $p = 0.088$ |

\* Statistically significant findings are highlighted for ease of reading.

**Figure 1.**
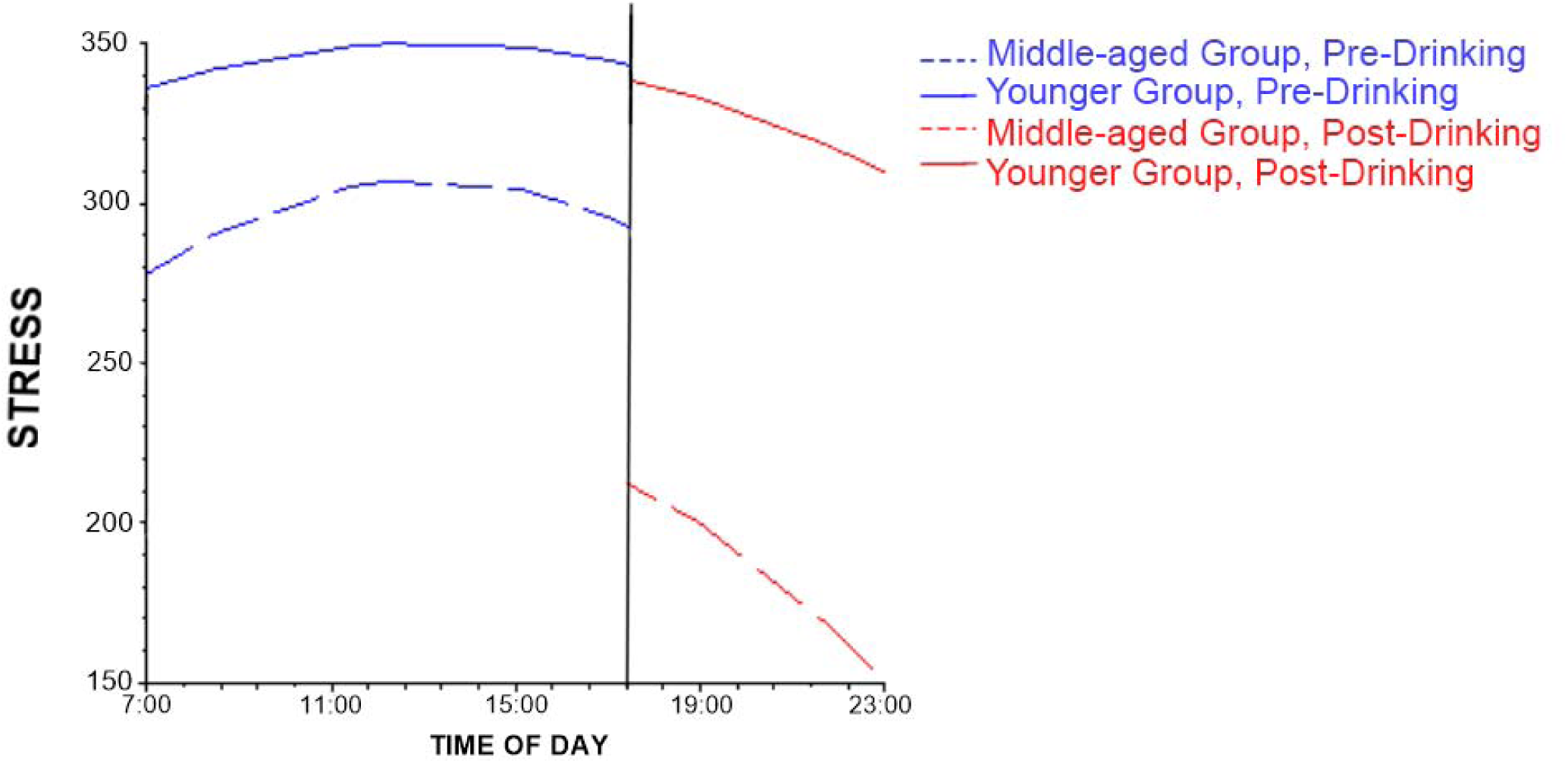
Visual representation of predicted stress ratings based on mixed-modeling results. Time of day is displayed on the x-axis, with a vertical line representing the timing of alcohol consumption. Predicted stress ratings are depicted on the y-axis. Dashed lines represent predicted responses from middle-aged adults, while solid lines represent predicted responses from younger adults; blue lines represent responses collected pre-drinking, while red lines represent responses collected post-drinking.

Analyses involving transformed outcome variables showed that although coefficient values and levels of statistical significance changed across analyses (as would be expected), overall patterns of effects remained the same across all iterations.

A post hoc analysis was conducted in an effort to determine if these results might be the result of different lifetime experiences. Scores on the Childhood Trauma Questionnaire [54-56], the Alcohol Craving Experience survey [57], the Multidimensional Assessment of Interoceptive Awareness [58], and the Body Perception Questionnaire short form [59], all collected during the initial screening visit, were compared across the groups,; however, no significant differences were observed.

## Discussion

The objective of the study was to examine daily stress and affective responses of younger and middle-aged adults who regularly consumed alcohol in response to brief periods of normal drinking and abstinence. We also assessed RSA_rest_ as a biomarker of stress. Based on existing evidence, we hypothesized that middle-aged adults would exhibit higher levels of stress across the day and poorer affect than younger adults, effects that would be corroborated by lower levels of RSA_rest_ and mitigated following episodes of drinking. The lack of difference in self-reported experience, both historical and current, suggests that the observed effects may be primarily associated with the measured biological features, namely age and autonomic function as measured via RSA_rest_.

Although we did observe that the middle-aged adults had lower RSA_rest_, contrary to our hypotheses and research in the general population [8-11,13], the middle-aged adults experienced higher affect throughout the day than the younger adults while stress levels were similar. Also, the stress levels of middle-aged adults actually decreased following drinking, whereas it was unchanged for the younger adults. It is possible that the patterns observed in the current study are sample specific; however, to our knowledge this is the first investigation using EMA methodology to examine stress and affect between younger- and middle-aged adults who habitually consume alcohol. Whereas further research on this population across the lifespan is warranted, habitual use of alcohol in moderation may function as an operable though transient coping mechanism for life stress [50,51,54], among middle-aged adults who have had a prolonged history of consumption. Of note, mounting evidence suggests deleterious health outcomes associated with daily, long-term alcohol consumption. If indeed further research corroborates that it reduces stress in middle-aged adults, this in situ stress relief must be counterbalanced against adverse effects such as increased risk of cancers (including breast, lip, and oral cavity cancers), ischemic heart disease, tuberculosis, and diabetes (GBD, 2018; Rumgay et al., 2021). Alternatively, long term, habitual use of alcohol may blunt or modify central processing of stress. As a first step, it would be informative to simply examine brain connectivity of younger and older adults to explore possible differences in neural networks that might offer a mechanism for such a hypothesis.

Another point that merits attention is the fact that the middle-aged adults who had lower levels of RSA_rest_, reported reduced stress compared to the younger adults. In the general population, RSA_rest_ has been found to be inversely related to stress ((Balzarotti et al., 2017; Beauchaine & Thayer, 2015; Fanning et al., 2020; Porges, 2007). Also, among males who are alcoholics, lower RSA_rest_ has been found to be associated with higher levels of stress and poor mood [46]. RSA_rest_ is a biomarker of stress that captures the consequences from multiple sources of insult across the lifespan, both physical and psychological (Porges, 2011b). In is important to underscore the fact the levels of RSA_rest_ are determined by both conscious and non-conscious processes, whereas self-report stress is a conscious process. Once again, this pattern in the data reinforce the importance of examining further whether or not habitual alcohol consumption into the middle years of life enhances well-being and served to buffer sources of insult that serve to lower RSA_rest_ [50,51,54].

There are limitations to this study that lay the foundation for further investigation. Primarily, this study has a small sample size, which needs to be expanded into larger samples before data can be extrapolated to the population. Secondly, this study recruited a very specific subset of drinkers; the results of these analyses may not replicate with those that would be observed in infrequent social drinkers or those with problems associated with their alcohol consumption. Since the psychological benefits observed in this study do not necessarily align with some expectations associated with autonomic focused research, future studies should also continue to explore differences in these patterns why accounting for HRV differences. However, even when considering these limitations, this study comprises an impactful finding for aging and alcohol research. These cross-sectional data suggest potential momentary stress relief to engaging with moderate alcohol consumption in a middle-aged population.

