## Supplement for "Differential momentary reports of stress and affect associated with alcohol consumption in middle-aged versus younger adults"

Hierarchical Modeling Equations:

Stress Level-1 Model:

Stress*_ij_* = β*_0j_* + β*_1j_**(Time of Day – Linear*_ij_*) + β*_2j_**(Time of Day – Quadratic*_ij_*) + β*_3j_**(Pre-Drinking*_ij_*) + β*_4j_**(Post-Drinking*_ij_*)

Stress Level-2 Model:

β*_0j_* = γ*_00_* + γ*_01_**(Middle-aged Group*_j_*) + μ*_0j_*

β*_1j_* = γ*_10_* + γ*_11_**(Middle-aged Group*_j_*)

β*_2j_*= γ*_20_* + γ*_21_**(Middle-aged Group*_j_*)

β*_3j_*= γ*_30_* + γ*_31_**(Middle-aged Group*_j_*)

β*_4j_*= γ*_40_* + γ*_41_**(Middle-aged Group*_j_*)

Stress Mixed Model:

Stress*_ij_* = γ*_00_* + γ*_01_**(Middle-aged Group*_j_*) + γ*_10_**(Time of Day – Linear*_ij_*) + γ*_11_**(Middle-aged Group*_j_*)*(Time of Day – Linear*_ij_*) + γ*_20_**(Time of Day – Quadratic*_ij_*) + γ*_21_**(Middle-aged Group*_j_*)*(Time of Day – Quadratic*_ij_*) + γ*_30_**(Pre-Drinking*_ij_*) + γ*_31_**(Middle-aged Group*_j_*)*(Pre-Drinking*_ij_*) + γ*_40_**(Post-Drinking*_ij_*) + γ*_41_**(Middle-aged Group*_j_*)*(Post-Drinking*_ij_*) + μ*_0j_* + *r_ij_*

Affect Level-1 Model:

Affect*_ij_* = β*_0j_* + β*_1j_**(Time of Day – Linear*_ij_*) + β*_2j_**(Time of Day – Quadratic*_ij_*) + β*_3j_**(Pre-Drinking*_ij_*) + β*_4j_**(Post-Drinking*_ij_*)

Affect Level-2 Model:

β*_0j_* = γ*_00_* + γ*_01_**(Middle-aged Group*_j_*) + μ*_0j_*

β*_1j_* = γ*_10_* + γ*_11_**(Middle-aged Group*_j_*)

β*_2j_*= γ*_20_* + γ*_21_**(Middle-aged Group*_j_*)

β*_3j_*= γ*_30_* + γ*_31_**(Middle-aged Group*_j_*)

β*_4j_*= γ*_40_* + γ*_41_**(Middle-aged Group*_j_*)

Affect Mixed Model:

Affect*_ij_* = γ*_00_* + γ*_01_**(Middle-aged Group*_j_*) + γ*_10_**(Time of Day – Linear*_ij_*) + γ*_11_**(Middle-aged Group*_j_*)*(Time of Day – Linear*_ij_*) + γ*_20_**(Time of Day – Quadratic*_ij_*) + γ*_21_**(Middle-aged Group*_j_*)*(Time of Day – Quadratic*_ij_*) + γ*_30_**(Pre-Drinking*_ij_*) + γ*_31_**(Middle-aged Group*_j_*)*(Pre-Drinking*_ij_*) + γ*_40_**(Post-Drinking*_ij_*) + γ*_41_**(Middle-aged Group*_j_*)*(Post-Drinking*_ij_*) + μ*_0j_* + *r_ij_*

**Table 4.** Stress modeling, adjusted for Total Drinking Counts^1^.

| **Intercept** |  |  |  |
| --- | --- | --- | --- |
| Overall | γ*_00_* = 365.62 | SE = 76.51 | *p* < 0.001 |
| Total Drink Count | γ*_01_* = 0.008 | SE = 0.03 | *p* = 0.977 |
| Middle-aged Group | γ*_02_* = -67.71 | SE = 52.51 | *p* = 0.206 |
| **Time of Day – Slope** |  |  |  |
| Overall | γ*_10_* = -2.99 | SE = 2.69 | *p* = 0.265 |
| Total Drink Count | γ*_11_* = 0.003 | SE = 0.01 | *p* = 0.741 |
| Middle-aged Group | γ*_12_* = -0.79 | SE = 1.79 | *p* = 0.661 |
| **Time of Day – Curvilinear** |  |  |  |
| Overall | γ*_20_* = -0.48 | SE = 0.47 | *p* = 0.314 |
| Total Drink Count | γ*_21_* = -0.005 | SE = 0.002 | *p* = 0.774 |
| Middle-aged Group | γ*_22_* = -0.74 | SE = 0.31 | *p* = 0.017 |
| **Pre-Drinking Slope** |  |  |  |
| Overall | γ*_30_* = 38.85 | SE = 28.98 | *p* = 0.169 |
| Total Drink Count | γ*_31_* = -0.28 | SE = 0.11 | *p* = 0.010 |
| Middle-aged Group | γ*_32_* = -31.57 | SE = 19.47 | *p* = 0.105 |
| **Post-Drinking Slope** |  |  |  |
| Overall | γ*_40_* = 52.09 | SE = 33.17 | *p* = 0.116 |
| Total Drink Count | γ*_41_* = -0.28 | SE = 0.12 | *p* = 0.017 |
| Middle-aged Group | γ*_42_* = -91.48 | SE = 22.77 | *p* < 0.001 |

^1^ Total Drink Count comes from the TLFB and is a self-reported count of participant’s consumption for the 90 days prior to their participation.

**^*^** Statistically significant findings are highlighted for ease of reading.

**Table 5.** Affect modeling, adjusted for Total Drinking Counts^1^.

| **Intercept** |  |  |  |
| --- | --- | --- | --- |
| Overall | γ*_00_* = 1.45 | SE = 0.66 | *p* = 0.036 |
| Total Drink Count | γ*_01_* = 0.002 | SE = 0.02 | *p* = 0.452 |
| Middle-aged Group | γ*_02_* = 1.26 | SE = 0.46 | *p* = 0.009 |
| **Time of Day – Slope** |  |  |  |
| Overall | γ*_10_* = 0.07 | SE = 0.02 | *p* = 0.002 |
| Total Drink Count | γ*_11_* = -0.00001 | SE = 0.00008 | *p* = 0.214 |
| Middle-aged Group | γ*_12_* = -0.05 | SE = 0.02 | *p* = 0.001 |
| **Time of Day – Curvilinear** |  |  |  |
| Overall | γ*_20_* = -0.005 | SE = 0.004 | *p* = 0.157 |
| Total Drink Count | γ*_21_* = -0.00001 | SE = 0.00001 | *p* = 0.402 |
| Middle-aged Group | γ*_22_* = -0.008 | SE = 0.002 | *p* = 0.002 |
| **Pre-Drinking Slope** |  |  |  |
| Overall | γ*_30_* = -0.12 | SE = 0.23 | *p* = 0.606 |
| Total Drink Count | γ*_31_* = 0.001 | SE = 0.001 | *p* = 0.366 |
| Middle-aged Group | γ*_32_* = -0.02 | SE = 0.15 | *p* = 0.873 |
| **Post-Drinking Slope** |  |  |  |
| Overall | γ*_40_* = 0.35 | SE = 0.26 | *p* = 0.175 |
| Total Drink Count | γ*_41_* = -0.001 | SE =0.001 | *p* = 0.259 |
| Middle-aged Group | γ*_42_* = 0.31 | SE = 0.18 | *p* = 0.082 |

^1^ Total Drink Count comes from the TLFB and is a self-reported count of participant’s consumption for the 90 days prior to their participation.

**^*^** Statistically significant findings are highlighted for ease of reading.

**Table 6.** Stress modeling, using stress z-scores^1^.

| **Intercept** |  |  |  |
| --- | --- | --- | --- |
| Overall | γ*_00_* = 0.18 | SE = 0.06 | *p* = 0.005 |
| Middle-aged Group | γ*_01_* = 0.21 | SE = 0.08 | *p* = 0.017 |
| **Time of Day – Slope** |  |  |  |
| Overall | γ*_10_* = -0.02 | SE = 0.008 | *p* = 0.023 |
| Middle-aged Group | γ*_11_* = 0.01 | SE = 0.01 | *p* = 0.367 |
| **Time of Day – Curvilinear** |  |  |  |
| Overall | γ*_20_* = -0.004 | SE = 0.001 | *p* = 0.006 |
| Middle-aged Group | γ*_21_* = -0.003 | SE = 0.002 | *p* = 0.129 |
| **Pre-Drinking Slope** |  |  |  |
| Overall | γ*_30_* = -0.17 | SE = 0.09 | *p* = 0.043 |
| Middle-aged Group | γ*_31_* = -0.13 | SE = 0.12 | *p* = 0.286 |
| **Post-Drinking Slope** |  |  |  |
| Overall | γ*_40_* = -0.11 | SE = 0.10 | *p* = 0.261 |
| Middle-aged Group | γ*_41_* = -0.47 | SE = 0.14 | *p* < 0.001 |

^1^ Z-scores were calculated for each participant as (rating – mean rating)/SD.

**^*^** Statistically significant findings are highlighted for ease of reading.

**Table 7.** Affect modeling, using affect z-scores^1^.

| **Intercept** |  |  |  |
| --- | --- | --- | --- |
| Overall | γ*_00_* = 0.13 | SE = 0.06 | *p* = 0.038 |
| Middle-aged Group | γ*_01_* = 0.08 | SE = 0.08 | *p* = 0.324 |
| **Time of Day – Slope** |  |  |  |
| Overall | γ*_10_* = 0.03 | SE = 0.01 | *p* < 0.001 |
| Middle-aged Group | γ*_11_* = -0.04 | SE = 0.01 | *p* < 0.001 |
| **Time of Day – Curvilinear** |  |  |  |
| Overall | γ*_20_* = -0.01 | SE = 0.001 | *p* < 0.001 |
| Middle-aged Group | γ*_21_* = -0.003 | SE = 0.002 | *p* = 0.077 |
| **Pre-Drinking Slope** |  |  |  |
| Overall | γ*_30_* = 0.07 | SE = 0.09 | *p* = 0.397 |
| Middle-aged Group | γ*_31_* = -0.04 | SE = 0.12 | *p* = 0.750 |
| **Post-Drinking Slope** |  |  |  |
| Overall | γ*_40_* = 0.11 | SE = 0.10 | *p* = 0.249 |
| Middle-aged Group | γ*_41_* = 0.17 | SE = 0.14 | *p* = 0.212 |

^1^ Z-scores were calculated for each participant as (rating – mean rating)/SD.

**^*^** Statistically significant findings are highlighted for ease of reading.

**Table 8.** Stress modeling, using Q5 stress scores^1^.

| **Intercept** |  |  |  |
| --- | --- | --- | --- |
| Overall | γ*_00_* = 0.31 | SE = 0.05 | *p* < 0.001 |
| Middle-aged Group | γ*_01_* = -0.004 | SE = 0.003 | *p* = 0.955 |
| **Time of Day – Slope** |  |  |  |
| Overall | γ*_10_* = 0.0004 | SE = 0.003 | *p* = 0.884 |
| Middle-aged Group | γ*_11_* = -0.01 | SE = 0.004 | *p* = 0.023 |
| **Time of Day – Curvilinear** |  |  |  |
| Overall | γ*_20_* = -0.001 | SE = 0.001 | *p* = 0.039 |
| Middle-aged Group | γ*_21_* = -0.001 | SE = 0.001 | *p* = 0.156 |
| **Pre-Drinking Slope** |  |  |  |
| Overall | γ*_30_* = -0.04 | SE = 0.03 | *p* = 0.194 |
| Middle-aged Group | γ*_01_* = -0.03 | SE = 0.05 | *p* = 0.512 |
| **Post-Drinking Slope** |  |  |  |
| Overall | γ*_40_* = -0.05 | SE = 0.04 | *p* = 0.168 |
| Middle-aged Group | γ*_41_* = -0.06 | SE = 0.05 | *p* = 0.246 |

^1^ Q5 scores are dichotomous, with participants categorized as reporting stress scores in the top quintile of responses, versus the rest of responders.

**^*^** Statistically significant findings are highlighted for ease of reading.

**Table 9.** Affect modeling, using Q5 affect scores^1^.

| **Intercept** |  |  |  |
| --- | --- | --- | --- |
| Overall | γ*_00_* = 0.15 | SE = 0.07 | *p* = 0.045 |
| Middle-aged Group | γ*_01_* = 0.32 | SE = 0.10 | *p* = 0.004 |
| **Time of Day – Slope** |  |  |  |
| Overall | γ*_10_* = 0.01 | SE = 0.002 | *p* = 0.037 |
| Middle-aged Group | γ*_11_* = -0.01 | SE = 0.004 | *p* < 0.001 |
| **Time of Day – Curvilinear** |  |  |  |
| Overall | γ*_20_* = -0.0004 | SE = 0.0004 | *p* = 0.334 |
| Middle-aged Group | γ*_21_* = -0.002 | SE = 0.001 | *p* = 0.001 |
| **Pre-Drinking Slope** |  |  |  |
| Overall | γ*_30_* = -0.01 | SE = 0.03 | *p* = 0.852 |
| Middle-aged Group | γ*_31_* = -0.01 | SE = 0.04 | *p* = 0.900 |
| **Post-Drinking Slope** |  |  |  |
| Overall | γ*_40_* = -0.06 | SE = 0.03 | *p* = 0.058 |
| Middle-aged Group | γ*_41_* = 0.15 | SE = 0.05 | *p* < 0.001 |

^1^ Q5 scores are dichotomous, with participants categorized as reporting affect scores in the top quintile of responses, versus the rest of responders.

**^*^** Statistically significant findings are highlighted for ease of reading.

**Table 10.** Stress modeling, using a Poisson distribution.

| **Intercept** |  |  |  |
| --- | --- | --- | --- |
| Overall | γ*_00_* = 1.34 | SE = 0.17 | *p* < 0.001 |
| Middle-aged Group | γ*_01_* = -0.24 | SE = 0.23 | *p* = 0.314 |
| **Time of Day – Slope** |  |  |  |
| Overall | γ*_10_* = -0.01 | SE = 0.01 | *p* = 0.070 |
| Middle-aged Group | γ*_11_* = -0.01 | SE = 0.01 | *p* = 0.501 |
| **Time of Day – Curvilinear** |  |  |  |
| Overall | γ*_20_* = -0.003 | SE = 0.001 | *p* = 0.011 |
| Middle-aged Group | γ*_21_* = -0.003 | SE = 0.002 | *p* = 0.043 |
| **Pre-Drinking Slope** |  |  |  |
| Overall | γ*_30_* = -0.15 | SE = 0.07 | *p* = 0.036 |
| Middle-aged Group | γ*_31_* = -0.07 | SE = 0.10 | *p* = 0.473 |
| **Post-Drinking Slope** |  |  |  |
| Overall | γ*_40_* = -0.07 | SE = 0.08 | *p* = 0.385 |
| Middle-aged Group | γ*_41_* = -0.35 | SE = 0.12 | *p* = 0.002 |

**^*^** Statistically significant findings are highlighted for ease of reading.
